# Anatomically distinct regions in the inferior frontal cortex are modulated by task and reading skill

**DOI:** 10.1101/2024.09.11.612349

**Authors:** Hannah L. Stone, Jamie L. Mitchell, Mia Fuentes-Jimenez, Jasmine E. Tran, Jason D. Yeatman, Maya Yablonski

## Abstract

Inferior frontal cortex (IFC) is a critical region for reading and language. This part of the cortex is highly heterogeneous in its structural and functional organization and shows high variability across individuals. Despite decades of research, the relationship between specific IFC regions and reading skill remains unclear. To shed light on the function of IFC in reading, we aim to (1) characterize the functional landscape of text-selective responses in the IFC, while accounting for interindividual variability; and (2) examine how text-selective regions in the IFC relate to reading proficiency. To this end, children with a wide range of reading ability (N=66; age 7-14 years, 34 female, 32 male) completed functional MRI scans while performing two tasks on text and non-text visual stimuli. Importantly, both tasks do not explicitly require reading, and can be performed on all visual stimuli. This design allows us to tease apart stimulus-driven responses from task-driven responses and examine where in the IFC task and stimulus interact. We were able to identify three anatomically-distinct, text-selective clusters of activation in the IFC, in the inferior frontal sulcus (IFS), and dorsal and ventral precentral gyrus (PrG). These three regions showed a strong task effect that was highly specific to text. Furthermore, text-selectivity in the IFS and dorsal PrG was associated with reading proficiency, such that better readers showed higher selectivity to text. These findings suggest that text-selective regions in the IFC are sensitive to both stimulus and task, and highlight the importance of this region for proficient reading.

**Significance statement:** The inferior frontal cortex (IFC) is a critical region for language processing, yet despite decades of research, its relationship with reading skill remains unclear. In a group of children with a wide range of reading skills, we were able to identify three anatomically distinct text-selective clusters of activation in the IFC. These regions showed a strong task effect that was highly selective to text. Text-selectivity was positively correlated with reading proficiency, such that better readers showed higher selectivity to text, even in tasks that did not require reading. These findings suggest that multiple text-selective regions within IFC are sensitive to both stimulus and task, and highlight the critical role of IFC for reading proficiency.

## Introduction

Inferior frontal cortex (IFC) has long been recognized as a critical region for language. This association dates back to the 19th century, with the discovery of an inferior frontal speech production region, now known as Broca’s area (Broca, 1861). By the 1990s, new high-resolution imaging methods enabled researchers to further parse functionally-specific subregions within IFC, including distinct regions for semantic and phonological processing (Shaywitz et al., 1994; Poldrack et al., 1999; Vigneau et al., 2006). Phonological processing, or the ability to manipulate the sounds of spoken language, is essential for fluent reading, and is often impaired in struggling readers (Bradley and Bryant, 1983; Wagner and Torgesen, 1987). Consequently, neuroimaging studies have focused on IFC as a region where differences between struggling and typical readers may emerge, yet these findings have been inconsistent (Norton et al., 2015). Some have reported reduced IFC activation in struggling readers compared to typical readers (Maisog et al., 2008; Zhang and Peng, 2022), while others found increased IFC activation in struggling readers (Pugh et al., 2001; Hoeft et al., 2007; Shaywitz and Shaywitz, 2008). Surprisingly, the most consistent location where group differences have been reported is not a phonological, but a visual region – the visual word form area (VWFA) in ventral occipitotemporal cortex (Kubota et al., 2019; Brem et al., 2020). Thus, despite the well-established role of IFC in phonological processing and language, its relationship with reading proficiency remains unclear.

The conflicting findings regarding the role of IFC in reading difficulties may arise from several factors, including variations in the tasks used (McNorgan et al., 2015; Norton et al., 2015), the stimulus and baseline conditions, participants’ age (Richlan et al., 2011), the region investigated within IFC, and lack of anatomical precision in defining regions across participants. Inter-subject variability can confound studies that transform individual functional data into a standard anatomical space and perform group-wise statistics. This “group average” approach can inadvertently group together different functional regions across participants, potentially averaging out meaningful differences (Glezer and Riesenhuber, 2013). This is especially pertinent in frontal cortex where anatomical and cytoarchitectural individual differences are prominent (Ruland et al., 2022; Nolan et al., 2024) and functional language regions have high anatomical variability (Fedorenko and Kanwisher, 2009).

An additional complexity of IFC is its role as a high-level regulatory region implicated in working memory, attention, and other high-level, domain-general processes (Miller and Cohen, 2001; Tops and Boksem, 2011). Several studies of vision have shown that activity in IFC is sensitive to both stimulus and task demands (McKee et al., 2014; Bugatus et al., 2017).

However, studies of reading in IFC have typically used language tasks, such as sentence reading, where teasing apart the stimulus-driven response from task demands is not straightforward. Thus, it remains unclear whether IFC is sensitive to visual text, or to the task demands imposed by reading, or their interaction.

To address these questions, here we aim to: (1) characterize the functional landscape of text-selective responses in IFC, while accounting for individual variability; (2) distinguish task effects, stimulus effects and task by stimulus interactions, and (3) examine how text-selective regions in IFC relate to reading skill. We adopted an individualized approach to identify text-selective regions on the cortical surface of children with a wide range of reading abilities. Importantly, our experimental paradigm manipulates task demands while keeping the stimuli and experimental structure constant, allowing us to tease apart stimulus-driven responses from task-driven responses. We use two tasks that do not explicitly require reading: a fixation task where participants make perceptual judgments on a fixation dot while viewing (but ignoring) images of different visual categories, and a one-back task, where they respond when the same image repeats. This task choice is critical to ensure that participants perform exactly the same task on both text (real words, pseudowords, consonant strings) and non-text (false fonts, faces, limbs, objects) stimuli and that individual differences are not confounded by differences in task performance.

## Methods

### Overview

This study is a subset of a larger, longitudinal study in which participants completed multiple behavioral assessments and MRI scans over the course of 13 months. Here we report on cross-sectional analyses from the baseline timepoint of the longitudinal study.

### Participants

A total of 78 subjects participated in the longitudinal study. For this analysis of the baseline timepoint, we excluded ten participants due to high motion in the scanner (for exclusion criteria see *BOLD response estimation* below) and two participants who did not complete the scan. Our final sample for analysis included 66 subjects (mean age 9.9 ± 1.37 years, range [7.7 -13.8], 34 female / 32 male) that had at least one usable run from each task. All subjects had normal or corrected-to-normal vision and no neurological disorders or hearing difficulty. All subjects were either monolingual English speakers or communicated in English at least 60% of the day, and had acquired English before the age of 3. Additionally, all subjects were screened using the Wechsler Abbreviated Scale of Intelligence, Second Edition (WASI-II; (Wechsler, 2011) to ensure their score was above the sixteenth percentile for their age group. All protocols were approved by the Stanford Internal Review Board on Human Subjects Research and informed assent and consent were obtained from each child and guardian, respectively.

### Behavioral assessments

Participants completed the Woodcock-Johnson Test of Achievement IV (Schrank and Wendling, 2018). The assessments were administered over Zoom as data collection for this study began during the COVID-19 pandemic. We used the Basic Reading Score (BRS) measure, which is an age-standardized composite of real word reading (letter-word ID subtest) and pseudoword reading (word attack subtest). Behavioral assessments took place within a couple of weeks of the MRI scan. Assessments were independently scored by two trained researchers and a third researcher was brought in to adjudicate any discrepancies.

### Experimental design

The experimental design followed the localizer experiment in (White et al., 2023) with some minor adjustments to make it suitable for children. We summarize the details here for convenience.

#### Stimuli

Subjects were presented with stimuli from 5 visual categories: real-letters, false-fonts, objects, faces, and limbs (see Figure 1). The real-letter category was composed of 4 sub-categories of strings: high-frequency words, low-frequency words, pronounceable pseudowords, and consonant strings. The false-font category included two different sets of false characters, one matched in low-level features to the Sloan font, and one matched in low-level features to the Courier font (Vidal et al., 2017). The stimuli were arranged such that a small image was presented at the center of the screen with a fixation dot (0.14 degree visual angle wide) and two larger images from the same visual category were presented on either side of fixation to balance the visual acuity and processing advantage of the fovea compared to the peripheral retina.

**Figure 1.**
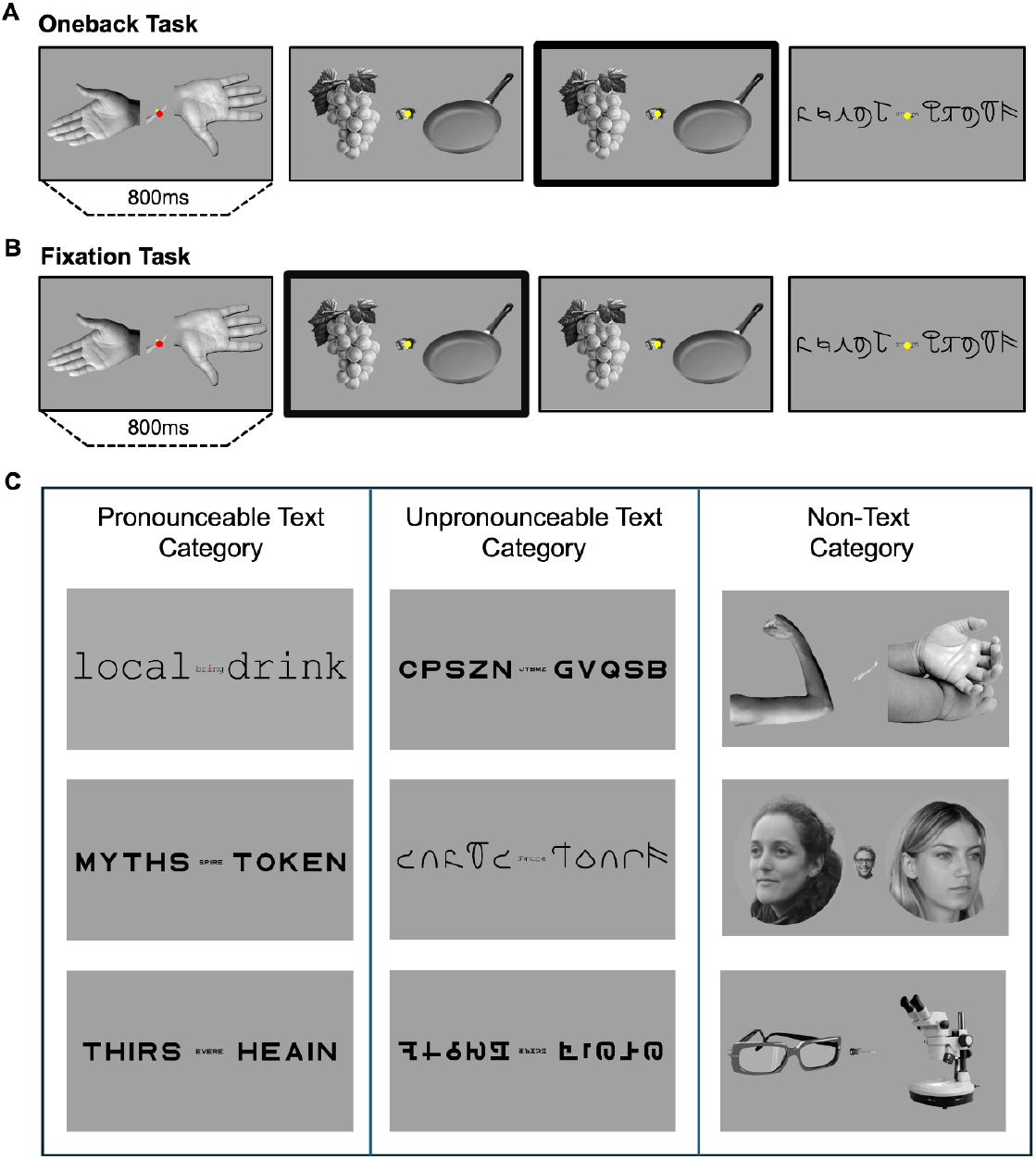
fMRI functional experiment. (A) One-back task: participants indicated via a button press when a stimulus appeared twice in a row. (B) Fixation task: participants indicated when a fixation dot at the center of the screen changed color. Stimuli were presented for 800ms followed by 200ms of a blank screen with a fixation dot. The frames where participants should respond in each task appear in bold. C) Example stimuli within three visual categories. The Pronounceable Text category included high and low frequency words and pseudowords, Unpronounceable Text included consonant strings and false-fonts, Non-Text included limbs, objects and faces. This illustration shows the faces of co-authors, but faces of people unfamiliar to participants were used in the actual experiment.

Each trial presented stimuli for 800 milliseconds followed by 200 milliseconds of a blank fixation screen. Four trials of a single visual category were presented in a row to form a block. A single run of the experiment randomly presented 10 blocks of each visual category, except for the real-letter category which included 20 blocks (5 blocks each of: high-frequency words, low-frequency words, pseudowords, and consonant strings).

#### Tasks

In each run, participants were asked to complete either a one-back task or a fixation task. In the one-back task, participants viewed the stimuli while fixating and responded via a button press when an image repeated twice in a row (Figure 1A). In the fixation task, participants viewed the stimuli while fixating and responded via a button press when the fixation dot changed color (Figure 1B). The one-back task was designed to draw the participant’s attention to the visual stimuli while the fixation task was designed to draw the participant’s attention away from the visual stimuli to the fixation dot. Image repetitions and fixation-dot color changes occurred at random in 33% of trials. Participants held the response box in their dominant hand and used their index finger to press the response button. Participants completed four runs of the experiment, alternating between the one-back task and the fixation task. Participants were visually monitored during each run and the entire run was discarded if the participant fell asleep at any point.

### MRI

#### Acquisition

Participants were scanned using a General Electric Sigma MR750 3T scanner at Stanford University’s Center for Cognitive and Neurobiological Imaging (CNI). Participants first completed an introduction session where they practiced a short version of the experiment in a scanner simulator to get acclimated to the scanner noises, the tasks, and the response box. During this session they also practiced staying still with a feedback system (MoTrak Head Motion Tracking System; RRID:SCR_009607).

Functional runs were acquired with a gradient echo EPI sequence using a multiband factor of 3 to achieve whole-brain coverage (51 slices). We used a TR of 1.19s with TE of 30ms and flip angle of 62 to obtain spatial resolution of 2.4 mm^3^ isotropic voxels. Each run included 232 frames and lasted 4:36 minutes. In addition, a high-resolution T1-weighted (T1w) anatomical scan was acquired with a spatial resolution of 0.9mm^3^ isotropic voxels.

#### Preprocessing

Preprocessing of functional data was performed using fMRIPrep 23.1.3 (Esteban et al., 2019), which is based on Nipype 1.8.6 (Gorgolewski et al., 2011, 2018).

#### Anatomical data preprocessing

The T1w image was reoriented into AC-PC alignment with ANTSpy version 0.4.2 (Tustison et al., 2021) and was used as an anatomical reference throughout the workflow. The T1w image was then skull-stripped using *Synthstrip* (Hoopes et al., 2022) and entered into Freesurfer’s recon-all pipeline (version 7.3.2; Dale et al., 1999) using *Synthseg* robust algorithm (Billot et al., 2023) for segmentation and surface reconstruction. All surfaces were visually inspected and edited manually in cases of inaccuracies. Freesurfer derivatives were then used by the fMRIPrep pipeline. Brain tissue segmentation of cerebrospinal fluid (CSF), white-matter (WM) and gray-matter (GM) was performed on the brain-extracted T1w using *fast* (FSL; Zhang et al., 2001). Volume-based spatial normalization to one standard space (MNI152NLin2009cAsym) was performed through nonlinear registration with *antsRegistration* (Tustison et al., 2021), using brain-extracted versions of both T1w reference and the T1w template.

#### Functional data preprocessing

For each BOLD run, a reference volume was generated using a custom methodology of fMRIPrep. Head-motion parameters with respect to the BOLD reference (transformation matrices and six rotation and translation parameters) were estimated before any spatiotemporal filtering using *mcflirt* (FSL; Jenkinson et al., 2002). A B0-nonuniformity map was estimated based on two echo-planar imaging (EPI) references with *topup* (Andersson et al., 2003). The estimated fieldmap was then aligned with rigid-registration to the target EPI reference run. The field coefficients were mapped on to the reference EPI using the transform. BOLD runs were slice-time corrected to the middle slice using *3dTshift* from AFNI (Cox and Hyde, 1997). The BOLD reference was then co-registered to the T1w reference using *bbregister* (FreeSurfer) which implements boundary-based registration (Greve and Fischl, 2009). The time-series of framewise motion parameters and the BOLD signal within the WM and the CSF were calculated and used later as confound factors in the BOLD response estimation. The BOLD time-series were resampled into standard space, generating a preprocessed BOLD run in MNI152NLin2009cAsym space. The BOLD time-series were also resampled onto the *fsnative* and *fsaverage* Freesurfer surfaces. All resamplings were performed with a single interpolation step by composing all the pertinent transformations (i.e. head-motion transform matrices, susceptibility distortion correction, and co-registrations to anatomical and output spaces).

Gridded (volumetric) resamplings were performed using *antsApplyTransforms* (ANTs), configured with Lanczos interpolation to minimize the smoothing effects of other kernels (Lanczos, 1964). Non-gridded (surface) resamplings were performed using *mri_vol2surf* (FreeSurfer).

#### BOLD response estimation

We ran *mriqc* (version 22.0.1; Esteban et al., 2017) to inspect the quality of the data and exclude noisy runs. Runs where the mean framewise displacement (FD) was 0.5mm or larger, and runs where more than 30% of frames had FD greater than 0.5mm, were excluded from analysis. In addition, runs where participants failed to keep their eyes open were also excluded from analysis. BOLD response estimation was performed by fitting a general linear model (GLM) using Nilearn (0.5.0; Nilearn contributors et al., 2024) to the BOLD time-series in each participant’s native surface. The design matrix for the GLM model included the white matter signal, the CSF signal and their first derivatives, as well as the 6 motion parameters. In addition, first and second order polynomial drift parameters were included in the design matrix. The estimated BOLD responses (beta weights) are expressed in units of percent signal change, and reflect the change in response relative to the blank trials when the participants were presented with a blank fixation screen.

#### Regions of Interest definition

We generated contrast maps comparing the estimated BOLD response to text stimuli (including high frequency and low frequency words, pseudowords, and consonant strings) with the responses to non-real letter stimuli (false fonts, faces, objects, limbs). Contrast maps were generated by weighing each visual category by the number of presented blocks to account for the higher number of text stimuli (20 blocks) compared with the other categories (10 blocks each). While we focus here on cross sectional data from a single timepoint, to increase SNR for delineating ROIs we created the contrast maps based on all available longitudinal data for each subject (6-20 runs, median=15, collapsing across both tasks). Contrast maps were thresholded at t-value > 2. Regions of interest (ROIs) were then defined for each individual in native space. The ROIs were defined with the following anatomical constraints which were defined based on each individual’s anatomy: the first activation cluster was identified in the inferior frontal sulcus (IFS). This ROI was defined between the anterior border of the IFS or by a vertical boundary line if the sulcus extended into the curvature of the prefrontal cortex, and the posterior border of the IFS. The IFS ROI was also constrained between the most dorsal sulcus of the IFS and the curvature of the inflated surface into the lateral sulcus. The next clusters of activation were identified in the precentral gyrus (PrG). This ROI was defined between the anterior edge of the precentral sulcus and the anterior edge of the central sulcus. Ventrally, the PrG ROI was constrained by the curvature of the inflated surface into the lateral sulcus and dorsally by a horizontal boundary line approximately ⅔ of the way up from the ventral boundary. Visual inspection revealed that activations within the PrG tended to cluster in two separate areas, thus we split this ROI into dorsal and ventral regions. To do this separation, a horizontal boundary line was defined on a group unthresholded probability map of the PrG. This boundary was projected into native space for each individual, effectively dividing their PrG ROI into dorsal and ventral components.

To investigate the relationship between reading skill and ROI size, a parallel set of ROIs were defined on contrast maps from the single time point (cross sectional) data only, using the same definition criteria. This was done to reduce the potential variability in ROI size associated with the varying number of runs included in the GLM.

#### Statistical analysis

We first extracted the response magnitude for each stimulus category and task in units of percent signal change (PSC) from the baseline (blank fixation screen), averaged within each ROI. We used linear mixed effect (LME) models as implemented in the *lme4* package in R (Bates et al., 2015) to evaluate how activation in the frontal lobe ROIs is influenced by stimulus category and task demands. For each ROI we fit a model with fixed effects of task (fixation vs. one-back) and stimulus category, where we grouped the stimuli into three qualitatively different categories: Pronounceable Text (high frequency and low frequency words, pseudowords), Unpronounceable Text (consonant strings and false-fonts), and Non-Text (faces, objects, limbs). We selected these groupings based on the phonological properties of the stimuli, reflecting the IFC’s role in the phonological processing of text. This approach was intended to highlight any potential differences in selective tuning to phonological features. All models included age and motion in the scanner as fixed effects, as well as random intercepts for each subject.

We next examined whether the ROIs exhibited similar response profiles. To this end, we fit a model with activation as the dependent variable and stimulus category and ROI as fixed variables (PSC ∼ Category*ROI + age + motion + (1 | subject)). We used sum coding for the ROI variable to avoid treating any specific ROI as the baseline. Instead, each ROI was compared to the grand mean. In addition, we fit a similar model where the dependent variable was the magnitude of the task effect, defined as the subtraction of fixation activation from one-back activation ((One-back -Fixation) ∼ Category*ROI + age + motion + (1 | subject)). This allowed us to first examine the response properties across ROIs over and above the different tasks, and then tease apart whether the task effect differs between the ROIs. We followed each significant interaction with post-hoc paired samples t-tests.

We then investigated whether reading level modulates activation in the frontal lobe. To this end we added z-scored individual reading scores (Basic Reading Score (BRS) from the Woodcock Johnson test) as a fixed effect to the models. For visualization, we split the sample into struggling readers and typical readers by a reading score cutoff of BRS<95. This cutoff equates to the 37th percentile as well as Woodcock-Johnson’s cut-off for ‘Average’ readers. In all statistical analyses, BRS score was treated as a continuous variable. We followed up on the ROI analysis with an exploratory vertex-wise analysis within a broad ROI encompassing the lateral frontal lobe, where we calculated the correlation between reading score and text selectivity (response to Pronounceable Text vs. all other categories) within each vertex. This complementary analysis provides a more detailed picture of the spatial organization of the relationship between reading skill and text-selectivity in the frontal lobe.

Lastly, to test whether reading skill was also associated with ROI size, we fit multiple linear regression models predicting ROI size (number of vertices) in the current timepoint from individual reading scores, age and motion.

## Results

Behaviorally, participants completed the one-back task with high accuracy (mean d’ = 1.72, 95% CI: [1.52, 1.92]) which was similar to their accuracy on the fixation task (mean d’ = 1.58, 95% CI: [1.42, 1.76]) (t = 1.304, p = 0.197). Generally participants responded faster in the one-back task with an average reaction time of 0.85 ± 0.24s compared with the fixation task at 0.92 ± 0.25s (t = -3.53, p = 0.001). Reading score (measured outside the scanner) was weakly correlated with accuracy in the one-back task (r = 0.255, p = 0.0386) but not in the fixation task (r = -0.025, p = 0.842).

### At least three anatomically distinct text selective regions in the frontal cortex

Text-selectivity was defined based on the contrast comparing responses to text stimuli (high frequency words, low frequency words, pseudo-words, consonant strings) with responses to all other categories (false-fonts, objects, faces, limbs). Visual inspection of these maps revealed at least three distinct activation peaks within each participant. Generally, the largest regions of activation were spread across the precentral gyrus (PrG) and clustered into dorsal and ventral regions. A more anterior patch (which was often diffuse and variable in size) was located along the inferior frontal sulcus (IFS). We identified and labeled these three text-selective ROIs within each participant by their anatomical location: IFS, dorsal PrG and ventral PrG (Figure 2). We used anatomical boundaries on each individual’s cortical surface to define these ROIs in order to maintain consistency across subjects (see *Methods*). We were able to identify all three ROIs in all 66 subjects. Figure 2 shows the spatial organization of these ROIs on the native cortical surface of four individual subjects, as well as probability maps showing the ROI locations on the *fsaverage* template. Even though the size and shape of these regions were variable across individuals, there was substantial overlap across the sample. However, in template space very few vertices contained the functional region in more than ⅓ of the participants, highlighting the importance of defining ROIs on the individual’s cortical surface.

**Figure 2.**
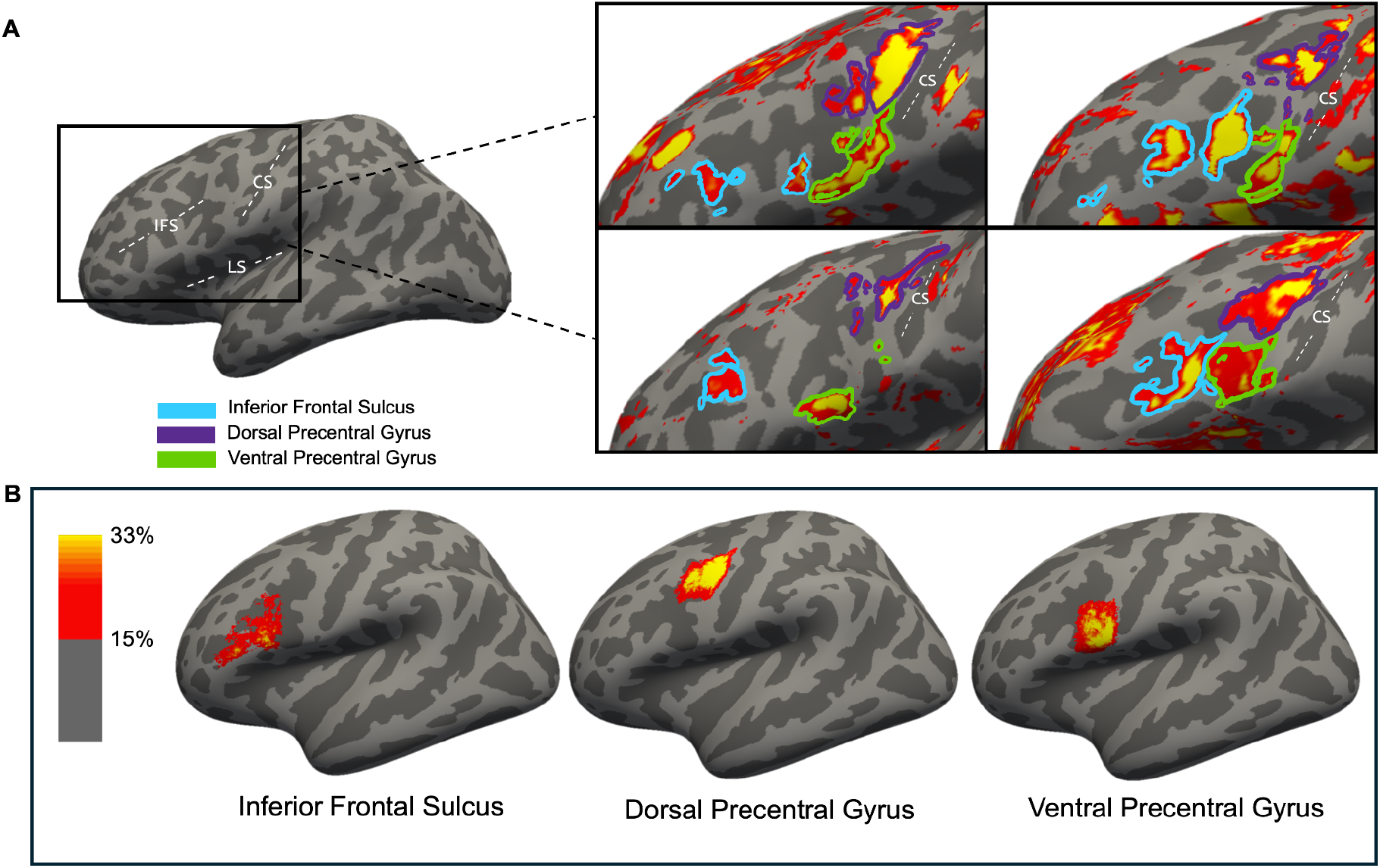
Anatomy of inferior frontal lobe text responses. (A) An example participant’s left hemisphere inflated surface with anatomical labels: IFS-inferior frontal sulcus, CS-central sulcus, LS-lateral sulcus. (B) Four example subject’s contrast maps, shown as heat maps (t-value > 2), in native space with hand-drawn ROI labels: inferior frontal sulcus ROI in blue, dorsal precentral gyrus ROI in purple, and ventral precentral gyrus ROI in green. Note that additional patches of frontal lobe text activation exist but we focused on the three regions that could be consistently defined in each individual. (C) Group average probability maps of ROI location in *fsaverage* space. Probability maps are thresholded at 15% overlap across subjects.

### Task enhances text activation in the frontal cortex

Across all ROIs, activation to Text stimuli was significantly higher than activation for the other categories, which is expected as this was the definition criterion for the ROIs. In fact, there was hardly any activation to non-text stimuli (Figure 3). In all three ROIs, activation for Pronounceable Text during the one-back task was significantly greater than during the fixation task (see Figure 3 and Table 1), suggesting that a stimulus-directed task evokes stronger neural response in these regions. Importantly, this task effect was highly specific for the Pronounceable Text stimuli, as indicated by a significant interaction between task and stimulus category (Table 1 and Figure 3). Specifically, the task effect was stronger in the Pronounceable Text category compared with the Non-Text category in all ROIs. This suggests that these regions are modulated by cognitive processes specific to text, as opposed to more domain-general mechanisms which would enhance activation of all attended visual stimuli. Interestingly, for the IFS alone, the task effect for Pronounceable Text was also significantly stronger than Unpronounceable Text, suggesting that the IFS is sensitive to the pronounceability of the stimuli.

**Table 1.** Linear mixed effect models examining the relationship between stimulus category (Pronounceable Text, Unpronounceable Text, and Non-Text) and task (one-back vs. fixation). Age and in-scanner motion (mean framewise displacement in mm) were entered as fixed effects of no interest. Non-Text included faces, limbs, and objects. UnProText (Unpronounceable Text categories) included consonant strings and false fonts. Pronounceable Text was defined as the reference level for the category variable, and fixation was the reference level for the task variable.

| Activation (PSC) ~ Category * Task + age + motion + (1 \| subject) |  |  |  |  |  |  |  |  |  |
| --- | --- | --- | --- | --- | --- | --- | --- | --- | --- |
|  | IFS |  |  | dorsal PrG |  |  | ventral PrG |  |  |
|  | β | t | p | β | t | p | β | t | p |
| (Intercept) | -0.209 | -0.857 | 0.3946 | 0.286 | 1.114 | 0.2695 | -0.043 | -0.160 | 0.8731 |
| Category |  |  |  |  |  |  |  |  |  |
| NonText | -0.182 | -6.980 | 0.0000 | -0.219 | -9.521 | 0.0000 | -0.241 | -8.696 | 0.0000 |
| UnProText | -0.131 | -4.507 | 0.0000 | -0.148 | -5.743 | 0.0000 | -0.128 | -4.117 | 0.0000 |
| Task |  |  |  |  |  |  |  |  |  |
| Oneback | 0.223 | 8.543 | 0.0000 | 0.240 | 10.407 | 0.0000 | 0.160 | 5.758 | 0.0000 |
| Category * Task |  |  |  |  |  |  |  |  |  |
| NonText : Oneback | -0.194 | -5.267 | 0.0000 | -0.166 | -5.106 | 0.0000 | -0.153 | -3.913 | 0.0001 |
| UnProText : Oneback | -0.138 | -3.355 | 0.0008 | -0.052 | -1.425 | 0.1546 | -0.027 | -0.606 | 0.5444 |
| Age | 0.032 | 1.521 | 0.1333 | -0.001 | -0.055 | 0.9562 | 0.014 | 0.595 | 0.5539 |
| Motion | 0.287 | 0.820 | 0.4151 | -0.557 | -1.511 | 0.1359 | 0.284 | 0.746 | 0.4584 |

**Figure 3.**
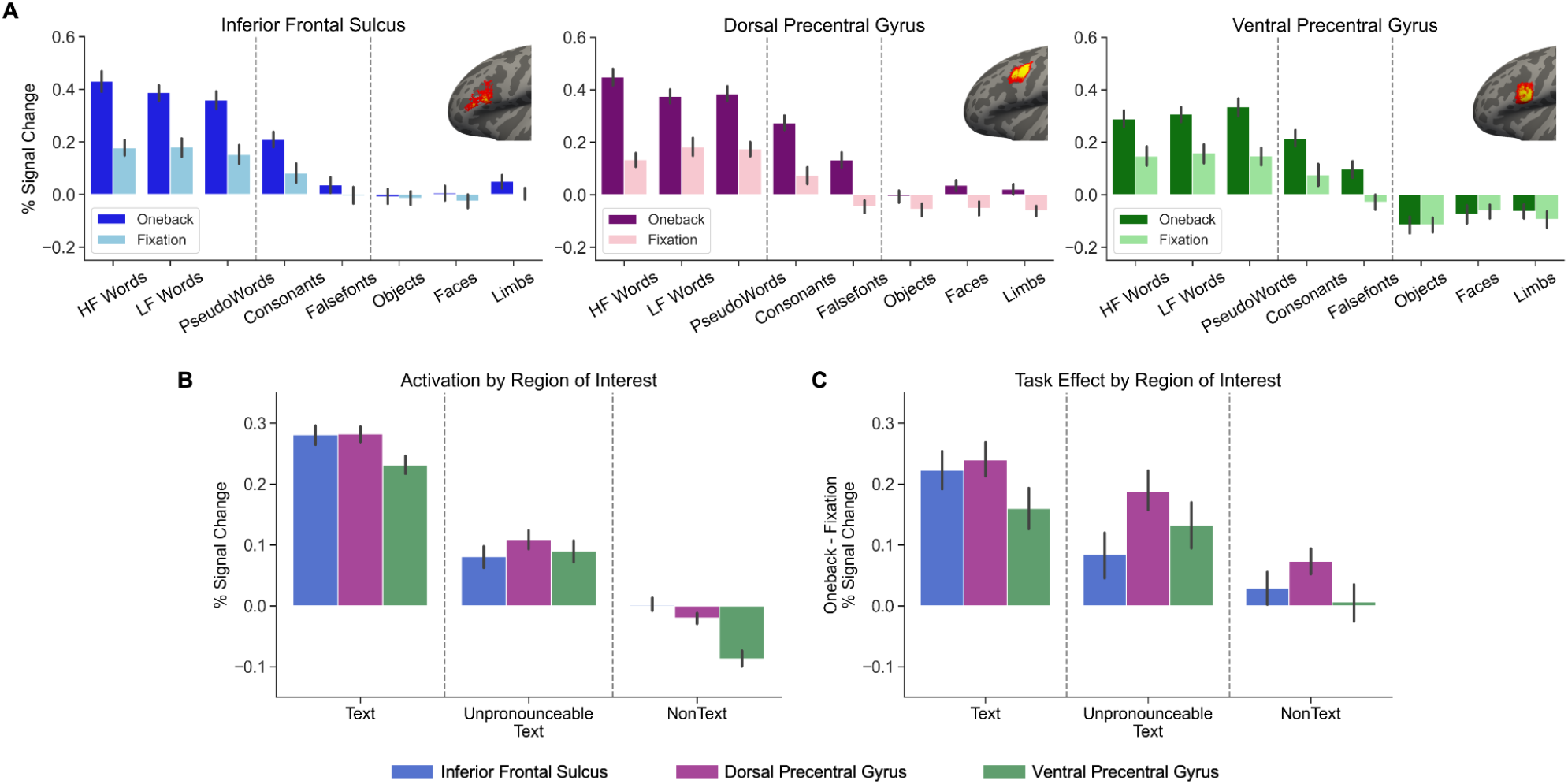
Individual ROI activation for each visual stimuli and task. (A) Activation is shown as mean percent signal change from baseline for each visual category and for each task (one-back and fixation). To represent the within subject variability, we subtracted each subject’s mean PSC from each datapoint and added the group mean. Error bars represent the 68% confidence interval (B) A comparison of the average activation between ROIs. Visual stimuli were grouped into text categories (Pronounceable Text, Unpronounceable Text, and Non-Text) for clarity. (C) The magnitude of task effect, calculated as the average difference between one-back and fixation PSC values within each subject.

To directly compare the ROIs we ran an additional LME where we evaluated whether the response magnitude was affected by the interaction between stimulus category and ROI (see *Methods* for model formula). There was no main effect of ROI (p > 0.1), indicating that all ROIs showed similar response magnitudes overall. There was also no interaction between ROI and stimulus category (p > 0.1), suggesting that the three regions show similar degrees of selectivity to text (Figure 3B). We next ran a similar LME predicting the magnitude of the task effect (as defined by the subtraction of activation during fixation from the activation during one-back).

There was no main effect of ROI on task effect magnitude (p > 0.05), indicating that the task effect was similar in strength across ROIs. Interestingly, we also observed a significant interaction between ROI and stimulus category (Figure 3C; Category (Unpronounceable Text) * ROI (IFS), β = -0.066, p = 0.0259). Post-hoc analyses revealed that the task effect was significantly stronger for Pronounceable Text compared with Unpronounceable Text in the IFS (paired t-test 3.896, p = 0.0002) while this was not the case in the dorsal PrG (t = 1.735, p = 0.086) or ventral PrG (t = 0.9, p = 0.369). This suggests that the IFS is sensitive to the pronounceability of the stimuli more than the PrG ROIs. Further, the PrG ROIs showed a significantly stronger task effect for Unpronounceable Text compared with Non-Text (t = 3.795, p = 0.0003; t = 3.882, p = 0.0002; dorsal and ventral respectively), while the IFS did not (t = 1.71, p = 0.092). This highlights the difference in response properties between the IFS and PrG ROIs.

### Reading skill modulates response to text in the frontal cortex

We next tested whether activation in each ROI was associated with reading skill. Combining across tasks, better reading skill was associated with stronger activation to Pronounceable Text, especially in the dorsal PrG (Table 2, Figure 4). In both the IFS and the dorsal PrG, the association between reading skill and activation strength was significantly stronger for Pronounceable Text stimuli, compared with Unpronounceable Text and with Non-Text stimuli

**Table 2.** Linear mixed effect models examining the relationship between reading skill and activation (top) or task effect (bottom). Task effect was defined as the difference between the one-back task and the fixation task.Reading was entered into the model as the z-scored Basic Reading Score (BRS) from the Woodcock-Johnson reading test. Age and in-scanner motion (mean framewise displacement in mm) were entered as fixed effects of no interest. NonText -non-textual visual categories (faces, limbs, objects). UnProText -unpronounceable text categories (consonant strings and false fonts). Pronounceable Text was defined as the reference level for the category variable.

| Activation (PSC) ~ Category * Reading + age + motion + (1 \| subject) |  |  |  |  |  |  |  |  |  |
| --- | --- | --- | --- | --- | --- | --- | --- | --- | --- |
|  | IFS |  |  | dorsal PrG |  |  | ventral PrG |  |  |
| | $\beta$ | t | p | $\beta$ | t | p | $\beta$ | t | p |
| (Intercept) | -0.103 | -0.422 | 0.6747 | 0.396 | 1.591 | 0.1168 | 0.029 | 0.110 | 0.9126 |
| **Category** |  |  |  |  |  |  |  |  |  |
| NonText | -0.279 | -14.612 | **0.0000** | -0.302 | -17.303 | **0.0000** | -0.318 | -15.841 | **0.0000** |
| UnProText | -0.201 | -9.391 | **0.0000** | -0.174 | -8.896 | **0.0000** | -0.141 | -6.282 | **0.0000** |
| **Reading (BRS z-score)** | 0.058 | 1.974 | 0.0518 | 0.092 | 3.107 | **0.0026** | 0.062 | 1.988 | 0.0502 |
| **Category \* Reading** |  |  |  |  |  |  |  |  |  |
| NonText : Reading | -0.048 | -2.512 | **0.0122** | -0.037 | -2.142 | **0.0325** | -0.020 | -0.977 | 0.3290 |
| UnProText : Reading | -0.043 | -1.996 | **0.0462** | -0.061 | -3.137 | **0.0018** | -0.005 | -0.223 | 0.8233 |
| Age | 0.032 | 1.510 | 0.1361 | -0.002 | -0.083 | 0.9339 | 0.013 | 0.585 | 0.5609 |
| Motion | 0.325 | 0.923 | 0.3595 | -0.478 | -1.330 | 0.1884 | 0.351 | 0.937 | 0.3525 |
| Task effect ~ Category * Reading + age + motion + (1 \| subject) |  |  |  |  |  |  |  |  |  |
|  | IFS |  |  | dorsal PrG |  |  | ventral PrG |  |  |
| | $\beta$ | t | p | $\beta$ | t | p | $\beta$ | t | p |
| (Intercept) | -0.203 | -0.558 | 0.5787 | -0.033 | -0.110 | 0.9128 | -0.060 | -0.145 | 0.8853 |
| **Category** |  |  |  |  |  |  |  |  |  |
| NonText | -0.194 | -7.257 | **0.0000** | -0.166 | -6.727 | **0.0000** | -0.153 | -5.583 | **0.0000** |
| UnProText | -0.138 | -4.623 | **0.0000** | -0.052 | -1.877 | 0.0612 | -0.027 | -0.865 | 0.3874 |
| **Reading (BRS z-score)** | 0.004 | 0.089 | 0.9291 | 0.036 | 0.974 | 0.3328 | -0.049 | -0.989 | 0.3258 |
| **Category \* Reading** |  |  |  |  |  |  |  |  |  |
| NonText : Reading | -0.035 | -1.324 | 0.1863 | 0.016 | 0.651 | 0.5155 | 0.064 | 2.331 | 0.0202 |
| UnProText : Reading | 0.015 | 0.497 | 0.6198 | 0.022 | 0.784 | 0.4336 | 0.035 | 1.130 | 0.2590 |
| Age | 0.034 | 1.064 | 0.2913 | 0.027 | 1.051 | 0.2972 | 0.026 | 0.720 | 0.4740 |
| Motion | 0.451 | 0.857 | 0.3946 | -0.006 | -0.014 | 0.9886 | -0.197 | -0.328 | 0.7437 |

**Figure 4.**
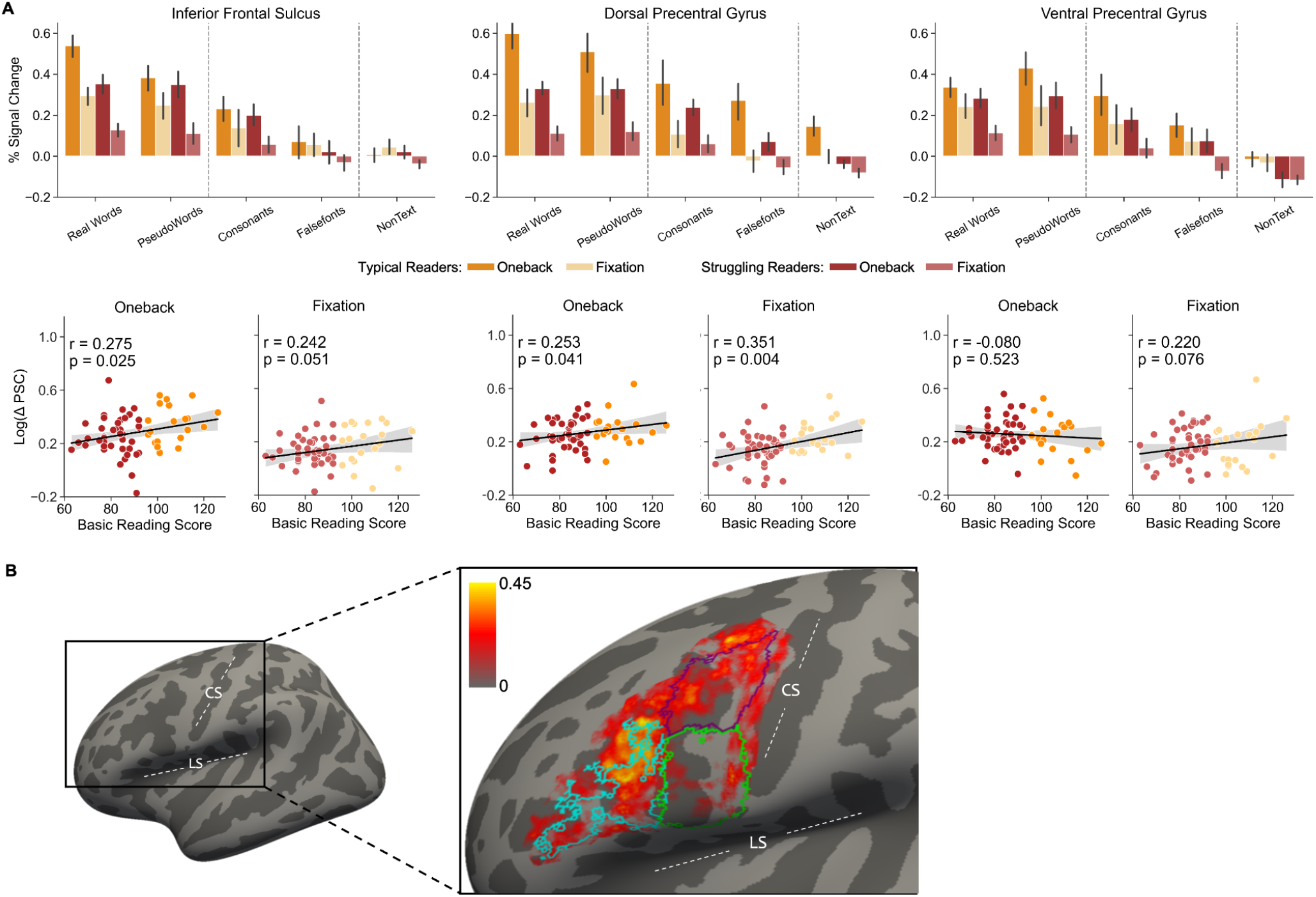
Relationship between activation in text-selective cortex and reading scores. (A) Average activation for each visual category, split by reading group and by task. Typical readers are indicated in orange (N=20), and struggling readers in red (N=46). The one-back task is indicated by the darker color and the fixation task by the lighter color within each reading group. Error bars represent a 68% confidence interval. Scatter plots underneath each ROI display the Pearson’s correlation between WJ Basic Reading Score and activation for Pronounceable Text -all other visual categories (log of Δ PSC), for each task. (B) Spatial distribution of the relationship between reading skill and text-selectivity. We transformed each participant’s z-scored contrast map from native surface to Freesurfer’s *fsaverage* template and calculated vertex-wise Pearson’s correlation between individual selectivity to text and WJ Basic Reading Score. Shown are the correlation coefficients for each vertex mapped onto an inflated *fsaverage* surface for visualization, overlayed with contours of the group average ROIs based on the group probability map (see Figure 2). The inferior frontal sulcus, dorsal precentral gyrus, and ventral precentral gyrus ROIs are labeled in blue, purple, and green, respectively. CS-central sulcus, LS-lateral sulcus.

(Table 2). This suggests that reading skill specifically modulates the response to Pronounceable Text in these regions. When focusing on the task effect as the predicted variable, reading skill was not associated with the magnitude of the task effect for text stimuli in any of the ROIs. In the ventral PrG alone there was a significant interaction such that reading skill was associated with larger task effect in the non-text category.

We followed up on the significant associations with reading by calculating post-hoc Pearson’s correlations between reading skill and text selectivity within each task separately (response to Pronounceable Text categories compared with all other categories) (Figure 4). This analysis showed that reading skill was positively correlated with text selectivity in the IFS (one-back r = 0.275, p = 0.025; fixation r = 0.242, p = 0.051) and the dorsal PrG (one-back r = 0.253, p = 0.041; fixation r = 0.351, p = 0.004), but less so in the ventral PrG (one-back r = -0.080, p = 0.523; fixation r = 0.220, p = 0.076). To take into account task performance, we repeated these analyses by fitting multiple regression models in each ROI, predicting text selectivity in each task from reading scores, task performance (d-prime), age and motion. This revealed that reading skill was a significant predictor of text selectivity in the IFS during the one-back task (β = 0.055, t = 2.253, p = 0.028) and in the dorsal PrG during the fixation task (β = 0.056, t = 3.014, p = 0.004). The rest of the predictor variables did not make a significant contribution to the models (p > 0.05) except for age, which was negatively associated with text selectivity during the fixation task in the ventral PrG alone (β = -0.037, t = -2.206, p = 0.031). This suggests that older children showed weaker text-selectivity in this specific ROI.

To further investigate the spatial distribution of the relationship between reading skill and text selectivity in the frontal lobe, we calculated the vertex-wise Pearson’s correlation between reading scores and text selectivity within a broad frontal lobe ROI. The resulting map (Figure 4B) shows high spatial overlap between the peak correlations and the dorsal parts of the IFS and dorsal PrG ROIs. This map shows a clear dissociation between the dorsal PrG and the ventral PrG, where the effect of reading skill is almost absent. This supports the notion that these ROIs are distinct and have different relationships with reading.

Finally, we tested whether reading skill was associated with ROI size. We found that struggling readers had smaller IFS ROIs (β = 9.892, p = 0.001, see Figure 5). However, the size of the dorsal PrG and ventral PrG ROIs was not associated with reading ability (β = 2.111, p > 0.1 and β = 1.145, p > 0.1, respectively). This again highlights the importance of the IFS region in reading, and the variability in the size of the region across participants. Age and motion were not associated with size in any of the ROIs (all p > 0.1).

**Figure 5.**
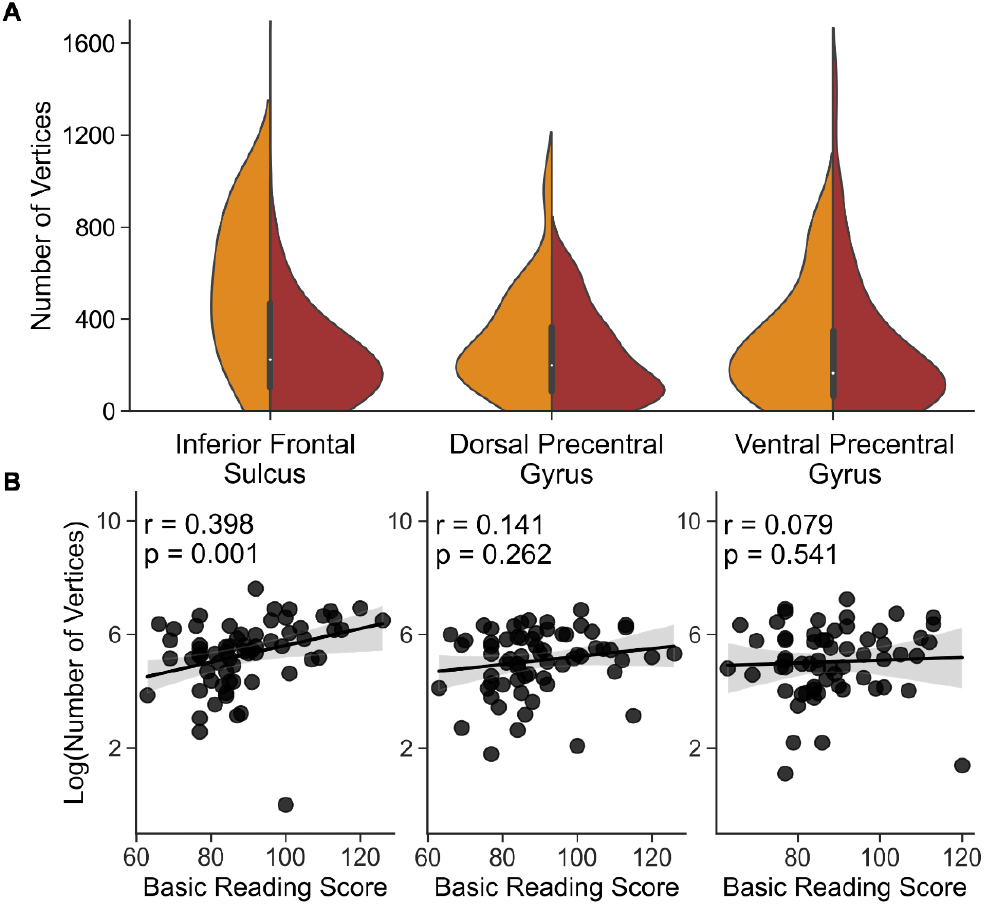
Reading skill is associated with ROI size. (A) Distribution of ROI size in number of vertices for typical readers (orange, N=20) and struggling readers (red, N=46). (B) Scatterplots display the Pearson’s correlation between the log number of vertices and WJ Basic Reading Score for each ROI.

## Discussion

Our findings indicate that three anatomically distinct, text-selective regions can be identified in the inferior frontal cortex of young readers. These regions show a dramatic effect of the cognitive task that the participant is engaged in, and this task effect is highly specific to text stimuli. Additionally, we found that individual differences in reading ability are related to activation levels in these regions, such that better readers show stronger activation for text stimuli, and larger clusters of text-selective responses in the IFS. Although all three regions were identified in the same way (based on their selectivity to text), our findings reveal distinct functional profiles: the IFS alone is sensitive to the pronounceability of presented text and shows a relationship with reading skill, the dorsal PrG shows a relationship with reading skill but is not sensitive to pronounceability, and the ventral PrG does not show a clear relationship with reading skill.

A growing number of studies have recognized that language regions in IFC comprise multiple, functionally distinct subregions (Fedorenko et al., 2012; Fedorenko and Blank, 2020; Wang et al., 2021, 2023; Chauhan et al., 2024). Some make the distinction between a phonological subregion, typically more posterior and dorsal, and a semantic subregion which is more anterior and ventral (Poldrack et al., 1999; Vigneau et al., 2006; Wang et al., 2021), in line with the dual-route model of language processing (Hickok and Poeppel, 2007; Saur et al., 2008). Specifically in the IFC, it has been suggested that the pars opercularis of “Broca’s area” plays a part in the dorsal, phonological, stream, while pars triangularis is part of the ventral, semantic, stream (Saur et al., 2008; Skeide et al., 2016). Others have noted that the broadly defined “Broca’s area” comprises a language subregion and a domain-general subregion that are spatially distinct (Fedorenko et al., 2012; Fedorenko and Blank, 2020). In the current study, we identified three different regions in IFC that are all highly selective to text, yet differ on several functional aspects. When considering the task effect, the more anterior IFS region shows the highest sensitivity to text pronounceability. In contrast, the dorsal and ventral PrG regions show a similar task effect to the text category and the unpronounceable text category. Additionally, both the IFS and the dorsal PrG show a strong relationship with reading skill, while this is not the case for the ventral PrG. The distinction between these ROIs was further confirmed by a vertex-wise analysis, which showed that peaks of correlation between text selectivity and reading skill overlapped with the IFS and the dorsal PrG, but not with the ventral PrG. This suggests that the anatomical distinctions we drew represent true functional differences between subregions of the IFC. We did not observe lexical or phonological effects in any of the ROIs (compare activation to high frequency words, low frequency words and pseudowords in Figure 3), perhaps due to the non-linguistic nature of the tasks which do not tap directly into lexical processes.

There is a growing understanding that task choice plays a key role in neuroimaging and has a critical influence on any measured response (Kay et al., 2023). A common practice to isolate language regions from other functional regions is to engage participants in a language task and compare it to a control task that does not recruit the language system. This approach manipulates both the stimulus category (e.g., words vs. faces) and the task (e.g., lexical decision vs. visual task) at the same time (e.g, Fedorenko and Blank, 2020; Borghesani et al., 2021; White et al., 2023). These comparisons are useful in identifying language regions, but conflate the stimulus-driven response with the cognitive task and the mental operations the participants perform on the stimulus. Thus to properly examine task effects in text-selective cortex it is crucial to hold the visual stimuli constant while varying the task, and to hold the task constant while varying the visual stimulus. In the visual system in particular, this approach showed that different tasks can enhance or attenuate the responses to different stimuli in category-selective cortex (Kay et al., 2015; Kay and Yeatman, 2017; White et al., 2023).

Interestingly, recent studies that examined task effects on a constant set of stimuli have found that patterns of neural representation in IFC were strongly task dependent, to a greater extent than in the ventral occipito-temporal cortex (Johnston and Everling, 2006; McKee et al., 2014; Bugatus et al., 2017; Jung and Walther, 2021). Further, McKee and colleagues (2014) found that in primates, tasks that recruited attention enhanced category selectivity in the IFC, in line with our current results. Along these lines, a recent study reported an interaction between task and stimulus in two subregions of IFC in adults (Chauhan et al., 2024). Our findings corroborate this observation and expand further by characterizing the spatial organization of text-selective regions in individual children, and uncovering their relationship with reading skill.

Another key point regarding task choice is our decision to use tasks that do not directly tap into language. Previous studies have typically located language related regions using tasks that explicitly require language processing, such as sentence or word reading, naming, lexical decision, or auditory phonological processing (Booth et al., 2007; Hoeft et al., 2007; Maisog et al., 2008; Fedorenko et al., 2012; Kovelman et al., 2012; Olulade et al., 2015; Wang et al., 2021, 2023; Zhang and Peng, 2022). When studying the relationship between brain activation and reading ability, this approach can intermingle individual differences with task difficulty: for example, in an extreme case, a participant that can’t read will not be able to complete a reading task and activation differences will be meaningless. A one-back task, on the other hand, can be done on any type of stimulus and task performance is orthogonal to reading ability. Comparing brain responses to a task that relies heavily on phonological processing will inevitably reveal group differences between children who struggle with the task itself and children who do not. It could be that the discrepancies between past studies reporting reduced or elevated IFC responses in struggling readers stem from employing tasks that differ on the extent to which they rely on phonological knowledge, reading, or language skills that are impaired in dyslexia. Here, we took special care to compare two tasks that do not explicitly require linguistic knowledge. Although we cannot fully rule out that some participants are automatically reading the text stimuli, it is not required in order to perform the task well. Our results therefore show that more skilled readers demonstrate higher tuning to visually presented text in IFC. Importantly, we find this effect even in the fixation task that directs attention away from the text/non-text stimuli.

The task effect we observed cannot be explained as a domain general, attention-driven amplification and, thus, we are careful not to use the term “attention” to describe the task effects reported here. Such a mechanism would result in increased activation in the one-back task for all visual stimuli (as studies have reported in the ventral occipitotemporal cortex (Kay and Yeatman, 2017), whereas we only see task-driven increased activation specific to text stimuli.

Additionally, we verify these regions are only selective for text rather than domain general in three ways: (1) the regions were defined on text-selective contrasts of real letters > all other visual stimuli, (2) we see hardly any activation for non-text categories (objects, faces, limbs), as opposed to category-selective regions in the occipitotemporal cortex that also respond to other visual categories to different degrees (Ben-Shachar et al., 2007; Centanni et al., 2017; Kay and Yeatman, 2017; Kubota et al., 2019; White et al., 2023; Dalski et al., 2024). (3) The task effect is absent for non-text categories, and differentiates between pronounceable and unpronounceable text in the case of the IFS. Therefore we conclude that this task-effect, or cognitive enhancement, is specific to text.

To conclude, we identified three anatomically distinct text-selective regions in IFC. These regions are highly selective to text even in tasks that do not explicitly require reading. We found that activation in these regions is associated with reading proficiency such that better readers show higher activation to text stimuli and greater text-selectivity. To our knowledge, this is the first study demonstrating that skilled readers have an increased response to purely visual text in the frontal cortex. However, it remains unclear how and when in the process of reading development this relationship comes about. The current cross-sectional data does not allow us to distinguish between two alternative hypotheses: It could be that accumulating reading experience tunes the responses in the frontal lobe to be more selective to text. Alternatively, it can be that children who develop more selective responses in the frontal cortex are the ones who will master reading more easily. Future studies employing causal and longitudinal methodologies will be needed to determine the existence and direction of a causal relationship between these regions and reading skill.

## Acknowledgements

This work was supported by NICHD R01-HD095861 to JDY. We thank the children and their families who contributed their time to participate in this study. We thank Megumi Takada, Clementine Chou and Kenny Tang for their invaluable part in data collection and Alex White for the task design.

The authors declare no competing financial interests.

